# SwiftOrtho: a Fast, Memory-Efficient, Multiple Genome Orthology Classifier

**DOI:** 10.1101/543223

**Authors:** Xiao Hu, Iddo Friedberg

## Abstract

**Introduction:** Gene homology type classification is a requisite for many types of genome analyses, including comparative genomics, phylogenetics, and protein function annotation. A large variety of tools have been developed to perform homology classification across genomes of different species. However, when applied to large genomic datasets, these tools require high memory and CPU usage, typically available only in costly computational clusters. To address this problem, we developed a new graph-based orthology analysis tool, SwiftOrtho, which is optimized for speed and memory usage when applied to large-scale data.

**Results:** In our tests, SwiftOrtho is the only tool that completed orthology analysis of 1,760 bacterial genomes on a computer with only 4GB RAM. Using various standard orthology datasets, we also show that SwiftOrtho has a high accuracy. SwiftOrtho enables the accurate comparative genomic analyses of thousands of genomes using low memory computers.

**Availability:** https://github.com/Rinoahu/SwiftOrtho

## Background

Gene homology type classification consists of identifying paralogs and orthologs across species. Orthologs are genes that evolved from a common ancestral gene following speciation, while paralogs are genes that are homologous due to duplication. Computationally detecting orthologs and paralogs across species is an important problem, as the evolutionary history of genes has implications for our understanding of gene function and evolution.

While the proper inference of homology type involves tracing gene history using phylogenetic trees [1], several proxy methods have been developed over the years. The most common method to infer orthologs by proxy is Reciprocal Best Hit or RBH [2, 3]. Briefly, RBH states the following: when two proteins that are encoded by two genes, each in a different genome, find each other as the best scoring match, they are considered to be orthologs [2, 3].

Inparanoid extends the RBH orthology relationship to include both orthologs and in-paralogs [4–6]. Specifically, Inparanoid distinguishes between orthologs and inparalogs, which were duplicated following a given speciation event [4–6]. It is then a matter of course to extend orthologous pairs between two species to an ortholog group, where an ortholog group is defined as a set of genes that are hypothesized to have descended from a common ancestor [6]. Several methods have been developed to identify ortholog groups across multiple species. These methods can be classified into two types: tree-based and graph-based. Tree-based methods construct a gene tree from an alignment of homologous sequences in different species and infer orthology relationships by reconciling the gene tree with its corresponding species tree [1, 7, 8]. Tree-based methods can infer a correct orthology relationship if the correct gene tree and species tree are given [9]. The main limitation of tree-based methods is the accuracy of the given gene tree and species tree. Erroneous trees lead to incorrect ortholog and in-paralog assignments [8–10]. Tree-based methods are also computationally expensive which limits the ability to apply them to large number of species [9, 11–13]. Graph-based methods infer orthologs and in-paralogs (Figure 1) from homologs and and then use different strategies to cluster them into orthologous groups [8, 11, 12]. The Clusters of Orthologous Groups or COG database detects triangles of RBHs in three different species and merges triangles with a common side [14]. Orthologous Matrix (OMA) clusters RBHs to orthologous groups by finding maximum weight cliques from the similarity graph [15]. MultiParanoid is an extension of Inparanoid, which uses InParanoid to detect triangle orthologs and in-paralogs in three different species as seeds and then merges the seeds into larger groups [16]. OrthoMCL also uses the InParanoid algorithm to detect orthologs, co-orthologs, and in-paralogs between two species [17] and then uses Markov Clustering (MCL) [18] to cluster these relationships into orthologous groups. Theoretically, graph-based methods are less accurate than tree-based methods, as the former identify orthologs and in-paralogs using proxy methods rather than directly inferring homology type from gene and species evolutionary history. In practice, graph-based methods have a similar accuracy as tree-based methods [9, 10, 19]. A comparison of several methods that include both tree-based and graph-based methods found that tree-based methods had even a worse performance than graph-based methods on large dataset [10]. One study compared several common methods including RBH, graph-based and tree-based and found that tree-based methods often give a higher specificity but lower sensitivity [20]. Several studies have also shown that graph-based methods find a better trade-off between specificity and sensitivity than tree-based methods [10, 20, 21]. Due to their better speed and accuracy, graph-based methods are generally preferred for analyzing large data set.

**Figure 1.**
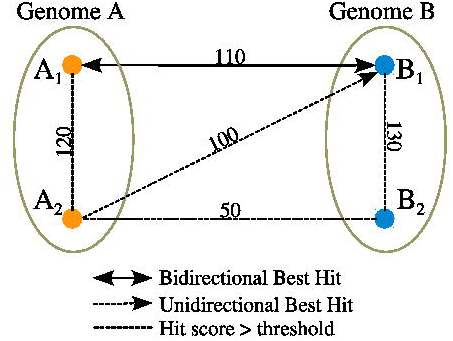
Orthology Inference Algorithm. Nodes are gene names, edges are similarity score of pairwise genes. 1. A_1_-B_1_ are putative orthologs identified by RBH. 2. A_1_-A_2_ and B_1_-B_2_ are putative in-paralogs as the bit scores of these pairs greater than A_1_-B_1_; 3. A_2_-B_1_ and A_2_-B_2_ are putative co-orthologs as these pairs are not orthologs but A_1_-B_1_ are orthologs and A_1_-A_2_, B_1_-B_2_ are in-paralogs.

Graph-based methods such as OrthoMCL and InParanoid can analyze hundreds of genomes, however they require considerable computational resources that may not be readily available [22, 23].

Here we developed a new orthology analysis tool named SwiftOrtho. SwiftOrtho is a graph-based method focused on speed, accuracy and memory efficiency. We compared SwiftOrtho with several existing graph-based tools using the gold standard dataset Orthobench [12], and the Quest for Orthologs service [24]. Using both benchmarks, we show that SwiftOrtho provides a high accuracy with lower CPU and memory usage than other graph-based methods.

## Methods

### Algorithms

SwiftOrtho is a graph-based orthology prediction method that performs homology search, orthology inference, and clustering by homology type.

### Homology Search

SwiftOrtho employs a seed-and-extension algorithm to find homologous gene pairs [25, 26]. At the seed phase, SwiftOrtho finds candidate target sequences that share common *k*-mers with the query sequence. *k*-mer size is an important factor that affects search sensitivity and speed [27, 28]. SwiftOrtho therefore uses long (≥ 6) *k*-mers to accelerate search speed. However, *k*-mer length is negatively correlated with sensitivity [27]. To compensate for the loss of sensitivity caused by increasing *k*-mer size, SwiftOrtho uses two approaches: non-consecutive *k*-mers and reduced amino-acid alphabets. Non-consecutive *k*-mer seeds (known as spaced seeds), were introduced in PatternHunter [17, 29]. The main difference between consecutive seeds and spaced seeds is that the latter allow mismatches in alignment. For example, the spaced seed `101101` allows mismatches at positions 2 and 5. The total number of matched positions in a spaced seed is known as a weight, so the weight of this seed is 4. A consecutive seed can be considered as a special case of spaced seed in which its weight equal its length. Spaced seeds often provide a better sensitivity than consecutive seeds [29, 30]. The default spaced seed patterns of SwiftOrtho are `1110100010001011, 11010110111` –two spaced seeds with weight of 8– but the user may define their own spaced seeds. Seed patterns were optimized using SpEED [30] and manual inspection. The choice of the spaced seeds and default alphabet are elaborated upon in the Methods section in the Supplementary Materials. At the extension phase, SwiftOrtho uses a variation of the Smith-Waterman algorithm [31], the *k*-banded Smith-Waterman or *k*-SWAT, which only allows for *k* gaps [32]. *k*-SWAT fills a band of cells along the main diagonal of the similarity score matrix (Figure 2B), and the complexity of *k*-swat is reduced to *O*(*k* ⋅ *min*(*n, m*)), where *k* is the maximum allowed number of gaps.

**Figure 2.**
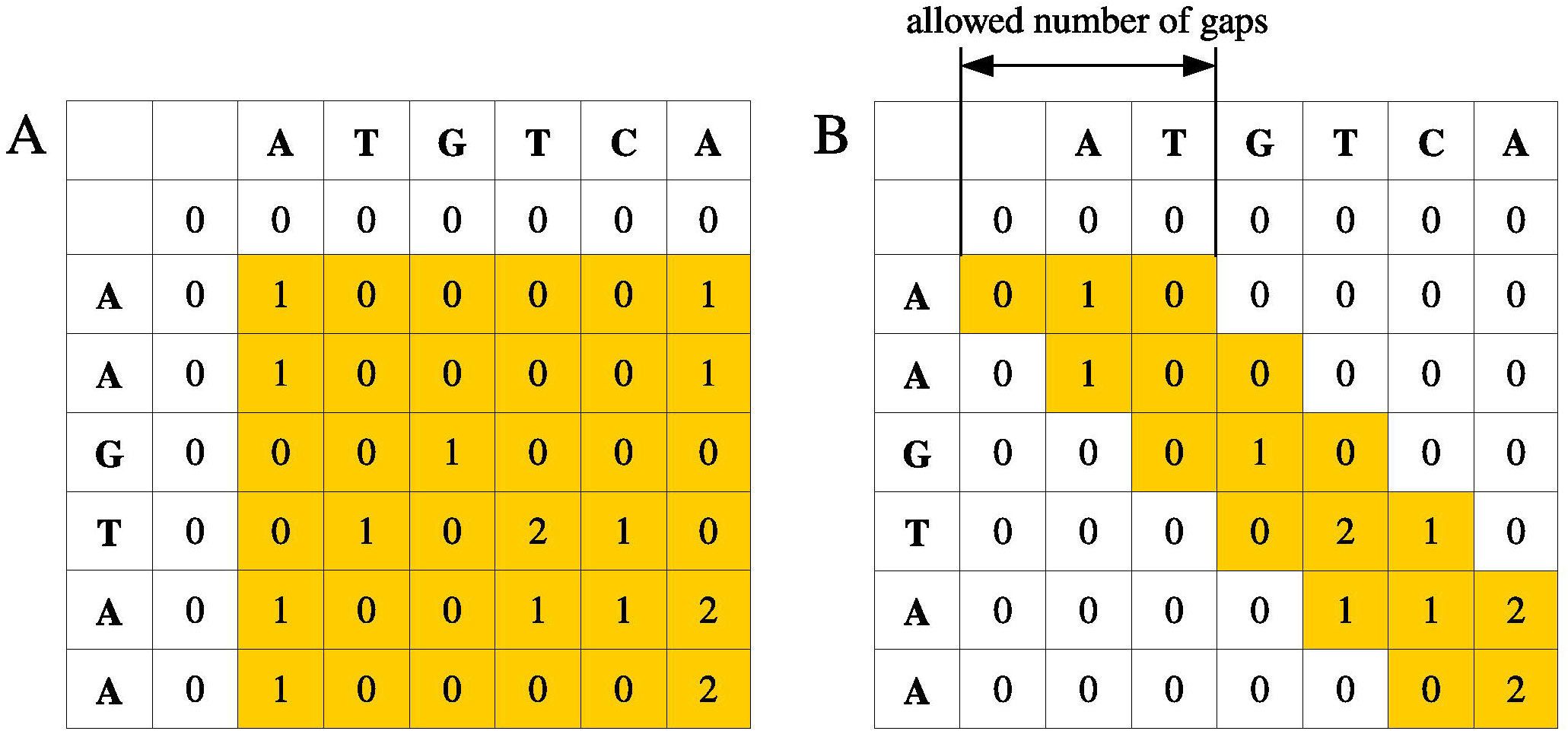
Comparing Standard Smith-Waterman with Banded Smith-Waterman. **A.** Similarity score matrix for Standard Smith-Waterman. Standard Smith-Waterman algorithm need to calculate all the entries. **B.** Similarity score matrix for Banded Smith-Waterman. Banded Smith-Waterman algorithm only need to calculate the entries on and near the diagonal.

Another method to mitigate the loss of sensitivity is to use reduced amino acid alphabets. Reduced alphabets are used to represent protein sequences using an alternative alphabet that combines several amino acids into a single representative letter, based on common physico-chemical traits [33–35]. Compared with the original alphabet of 20 amino acids, reduced alphabets usually improve sensitivity [36, 37]. However, reduced alphabets also introduces less specific seeds than the original alphabet, which reduces the search speed.

### Orthology Inference

SwiftOrtho employs a graph-based approach as the method to infer orthologs, co-orthologs and in-paralogs from homologs (Figure 1), and uses RBH to identify the orthologs. If the bit score between gene A_1_ and A_2_ in genome A is higher than that between A_1_ and all its orthologs in other genomes, A_1_ and A_2_ are considered in-paralogs in genome A. if A_1_ in genome A and B_1_ in genome B are orthologs, in-paralogs of A_1_ and B_1_ are co-orthologs (Figure 1). This process requires many queries so it is therefore better to store the data in a way that facilitates fast querying. SwiftOrtho sorts the data and uses a binary search algorithm to query the sorted data, which significantly reduces memory usage when compared with an Relational Database Management System or a hash table. With the help of this query system, SwiftOrtho can process data that are much larger than the computer memory.

After inferring orthology, the inferred orthology relationships are treated as the edges of a graph. Each edge is assigned a weight for cluster analysis. Appropriate edge-weighting metrics can improve the accuracy of cluster analysis. Gibbons compared the performance of several BLAST-based edge-weighting metrics and uses the bit score [38]. SwiftOrtho also uses the normalized bit score as edge-weighting metric. The normalization step take the same approach as OrthoMCL [22]: For orthologs or co-orthologs, the weight of (co-)ortholog (Figure 1) A_1_ in genome A and B_1_ in genome B is divided by the average edge-weight of all the (co-)orthologs between genome A and genome B. For in-paralogs, SwiftOrtho identifies a subset S of all in-paralogs in genome A, with each in-paralog A_*x*_-A_*y*_ in subset S, A_*x*_ or A_*y*_ having at least one ortholog in another genome. The weight of each in-paralog in genome A is divided by the average edge-weight of subset S in genome A [22].

### Clustering Orthology Relationships into Orthologous Groups

SwiftOrtho provides two methods to cluster orthology relationships in orthologous groups. One is the Markov Cluster algorithm (MCL), an unsupervised clustering algorithm based on simulation of flow in graphs [18]. MCL is fast and robust on small networks and has been used by several graph-based tools [17, 39–41]. However, MCL may run out of memory when applied on a large-scale network. To reduce memory usage, we cluster each individual connected component instead of the whole network because there is no flow among components [18]. However, for large and dense networks a single connected component could still be too large to be loaded into memory.

For the large networks, SwiftOrtho uses an Affinity Propagation Clustering algorithm (APC)[42]. The APC algorithm finds a set of centers in a network, where the centers are the actual data points and are called “exemplars”. To find exemplars, APC needs to keep two matrices of the responsibility matrix *R* and the availability matrix *A*. The element *R_i,k_* in *R* reflects how well-suited node k is to serve as the exemplar for node i while the element *A_i,k_* in *R* reflects how appropriate node i to choose node k as its exemplar [42]. APC uses Equation 1 to update *R*, and Equation 2 to update *A*, where *i, k, i*^′^*, k*^′^ denote the node number, and *S_i,k_*^′^ denotes the similarity between node *i* and node *k*^′^.

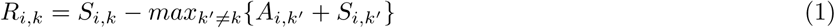

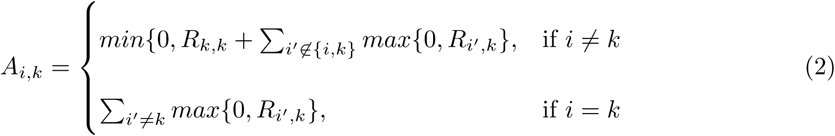

The node *k* that maximizes *A_i,k_* + *R_i,k_* is the exemplar of node *i*, and each node *i* is assigned to its nearest exemplar. APC can update each element of matrix *R*and *A* one by one, so, it is unnecessary to keep the whole matrix of *R* and *A* in memory. Generally, the time complexity of APC is O(*N* ^2^ ⋅ *T*) where *N* is number of nodes and *T* is number of iterations [42]. In this case, the time complexity is *O*(*E* ⋅ *T*), where *E* stands for edges which is number of orthology relationships and *T* is number of iterations. We implemented APC in Python, using Numba [43] to accelerate the numeric-intensive calculation parts.

## Application to Real data

### Data Sets

We applied SwiftOrtho to three data sets to evaluate its predictive quality and performance:

1. The *Euk* set was used to evaluate the quality of predicted orthologous groups. This set contains 420,415 protein sequences from 12 eukaryotic species, including *Caenorhabditis elegans, Drosophila melanogaster, Ciona intestinalis, Danio rerio, Tetraodon nigroviridis, Gallus gallus, Monodelphis domestica, Mus musculus, Rattus norvegicus, Canis familiaris, Pan troglodytes* and *Homo sapiens*. The protein sequences for these genes were downloaded from EMBL v65 [44].
2. The *QfO 2011* set was used to evaluate the quality of predicted orthology relationships. This set was the reference proteome dataset (2011) of The Quest for Orthologs[24], which contains 754,149 protein sequences of 66 species.
3. The large *Bac* set was used to evaluate performance, including CPU time, real time and RAM usage. This set includes 5,950,817 protein sequences from 1,760 bacterial species. The protein sequences were downloaded from GenBank [45]. For a full list, see the additional file 1.

### Comparing SwiftOrtho with existing Tools

We compared SwiftOrtho with several existing orthology analysis tools for predictive quality and performance. The methods compared were: OrthoMCL(v2.0), FastOrtho, OrthAgogue, and OrthoFinder.

### Orthology Analysis Pipeline

The pipeline for all the tools follows the standard steps of graph-based orthology prediction, (1) all-*vs*-all homology search, (2) orthology inference, and (3) cluster analysis.

#### Homology Search

SwiftOrtho used its built-in module to perform all-*vs*-all homology search. For all the three sets, the E-value was set 10^−5^. The amino acid alphabet was set to the regular 20 amino acids for the three sets. The spaced seed parameter was set to `1011111,11111` for the Euk, `11111111` for the *QfO 2011*, and `111111` for *Bac*.

OrthoMCL, FastOrtho, OrthAgogue, and OrthoFinder used BLASTP (v2.2.26) to perform all-*vs*-all homology search. The first three tools require the user to do this manually. In order to be able to compare, the −e (e-value), −v (number of database sequences to show one-line descriptions), and −b (number of database sequence to show alignments) parameters of BLASTP were set to 10^−5^, 1,000,000, and, 1,000,000 for OrthoMCL, FastOrtho, and OrthAgogue. The OrthoFinder calls BLASTP, and the E-value of BLASTP have been set to 10^−3^.

#### Orthology Inference

SwiftOrtho, OrthoMCL, FastOrtho, OrthAgogue, and OrthoFinder were applied to perform orthology inference on the homologs. The first four tools are able to identify (co-)orthologs and in-paralogs, and the coverage (fraction of aligned regions) was set to 50%, while other parameters were set to their default values, see Supplementary Materials for full details. FastOrtho does not report (co-)orthologs and in-paralogs directly. However, the relevant information is stored in an intermediate file, from which we have extracted that information. Orthofinder does not report orthology relationships.

#### Cluster Analysis

All the tools in this study use MCL [18] for clustering. To control the granularity of the clustering, MCL performs an inflation operation controlled by *-I* option [18, 46]. In this study, *-I* was set to 1.5. To take advantage of multiprocessor capabilities, we set the thread number of MCL to 12. SwiftOrtho has an alternative clustering algorithm APC, which we have also applied to *Euk* and*Bac*.

### Evaluation of Prediction Quality

#### Evaluation of Predicted Orthologous Group

The OrthoBench set was used to evaluate the quality of predicted orthologous groups in *Bac*. This set contains 70 manually curated orthologous groups of the 12 species from *Bac* and has been used as a high quality gold standard benchmark set for orthologous group prediction [12], we used OrthoBench v2 (Supplementary Table S1). In this study, each manually curated group of OrthoBench v2 set finds the best match in the predicted orthologous groups, where the best match means that the number of genes shared between manually curated and predicted orthologs is maximized, and the precision and recall are calculated(Figure 3A.).

**Figure 3.**
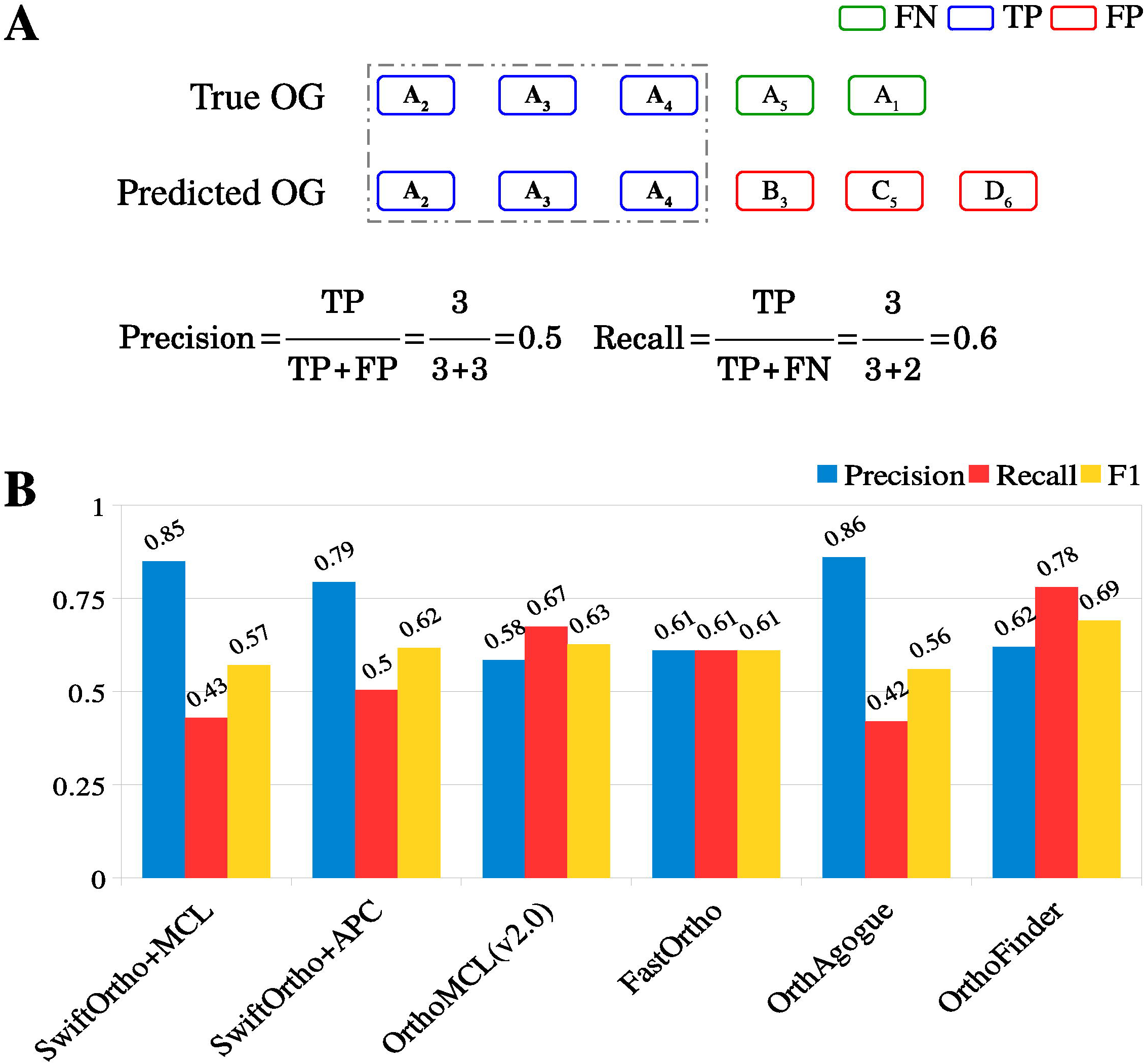
Evaluation of predicted orthologous groups. **A.** Definition of precision and recall. **OG:** orthologous group, **FN:** genes only found in true orthologous group,**TP:** genes shared between true and predicted orthologous group, **FP:** genes only found in predicted orthologous group; B. Evaluation of different tools on OrthoBench2. **SwiftOrtho+MCL:** SwiftOrtho with MCL; **SwiftOrtho+APC:** SwiftOrtho with Affinity Propagation Clustering.

#### Evaluation of Predicted Orthology Relationships

The *Quest of Orthologs* web-based service (QfO) was employed to evaluate the quality of the orthology relationships predicted from the *QfO 2011* set[24]. QfO service evaluates the predictive quality by performing four phylogeny-based tests of *Species Tree Discordance Benchmark*, *Generalized Species Tree Discordance Benchmark*, *Agreement with Reference Gene Phylogenies: SwissTree*, and *Agreement with Reference Gene Phylogenies: TreeFam-A*, and two function-based tests of *Gene Ontology conservation test* and *Enzyme Classification conservation test* [24]. We also applied two more orthology prediction tools, SonicParanoid[47] and InParanoid (v4.1)[4], on the *QfO 2011* set and used their results as control. The pairwise orthology relationships were extracted from the predicted orthologous groups of all the tools, including SonicParanoid and InParanoid, and then submitted to the QfO web-service for further evaluation.

### Hardware

Unless specified otherwise, all tests were run on the Condo cluster of Iowa State University with Intel Xeon E5-2640 v3 at 2.60GHz, 128GB RAM, 28TB free disk. The Linux command */usr/bin/time -v* was used to track CPU and peak memory usage.

## Results

We compared the orthology analysis performance of SwiftOrtho, OrthoMCL, FastOrtho, OrthAgogue, and OrthFinder using *Euk*, *QfO 2011*, and *Bac*. The orthology analysis consists of homology search, orthology inference, and cluster analysis.

### Orthology Analysis on *Euk*

The results of orthology analysis on *Euk* are summarized in Table 1:

**Table 1.**
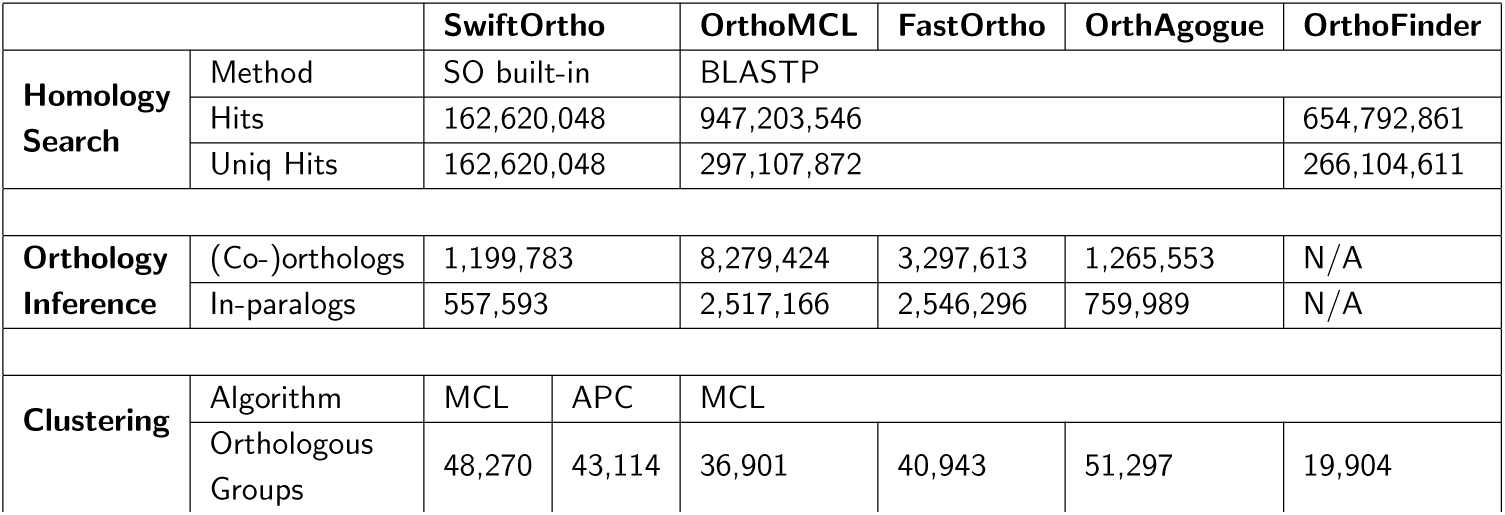
Comparative orthology analysis on the Euk set. N/A: not available, SO: SwiftOrtho, MCL: Markov Clustering, APC: Affinity Propagation Cluster.

### Homology Search

The homology search results show that BLASTP detected the largest number of homologs (947,203,546). SwiftOrtho found 57.5% of the homologs detected by BLASTP but was 38.7 times faster than BLASTP. SwiftOrtho used longer *k*-mers, which reduced both specific and non-specific seed extension. The longer *k*-mers cause seed-and-extension methods to ignore low similarity sequences. According to the RBH rule, orthologs should have higher similarity than non-orthologs, so, the decrease in homolgs of SwiftOrtho does not significantly affect the next orthology inference. We compared RBHs inferred from homologs detected by BLASTP and SwiftOrtho, and the numbers of RBHs for BLASTP and SwiftOrtho are 654,730 and 645,091, respectively. Identical RBHs are 497,286 (76.0% of BLASTP). These results shows that although SwiftOrtho found fewer homologs than BLASTP, SwiftOrtho does not significantly reduce the number of RBHs. The following results in Figure 4 also show that there is no significant difference between SwiftOrtho and BLASTP in orthologous groups prediction.

**Figure 4.**
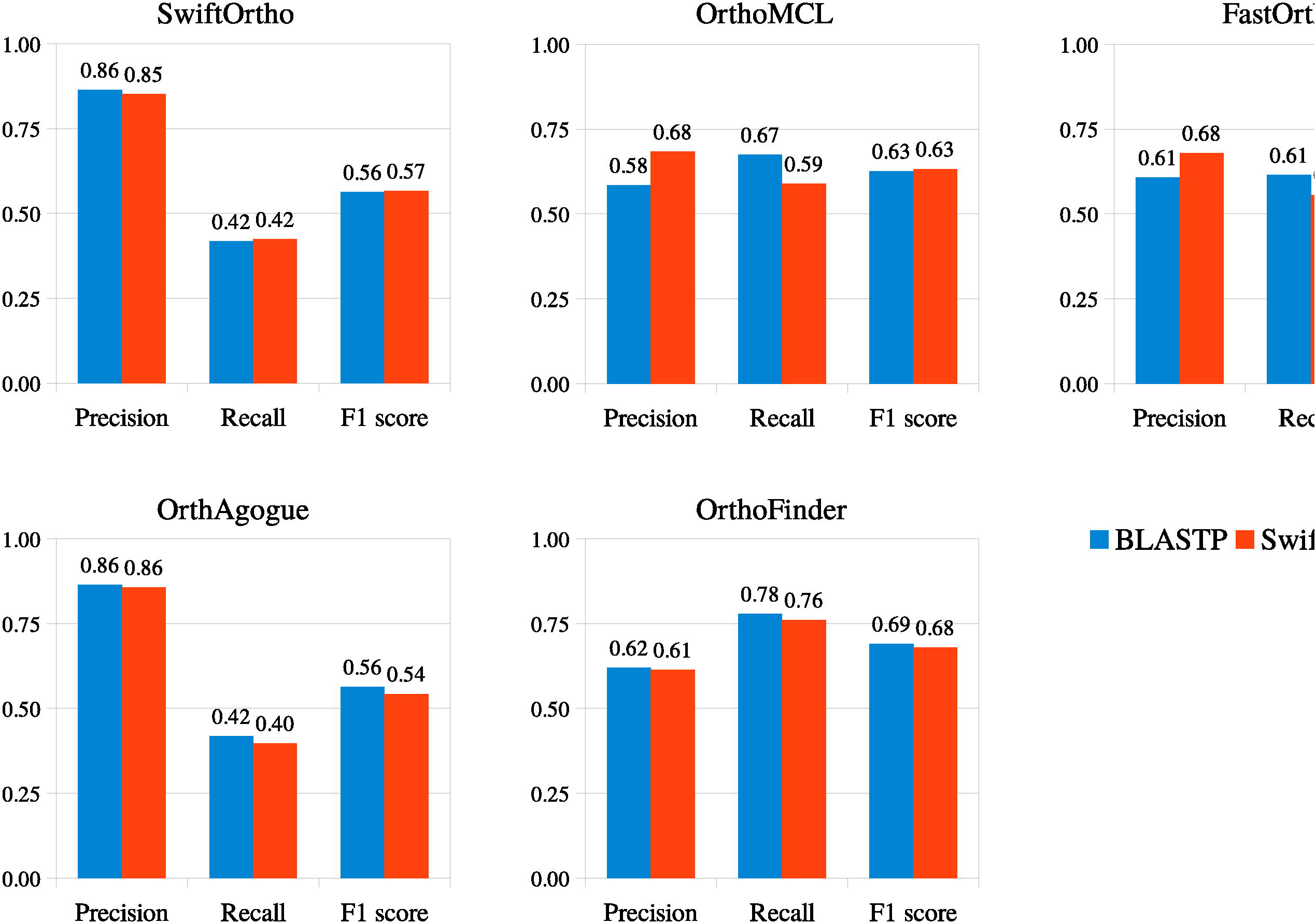
Comparing BLASTP and SwiftOrtho’s homology search module on the quality of orthologous groups prediction. BLASTP and SwiftOrtho’s search module perform an all-*vs*-all search on the *Euk* set, respectively. Then, all the orthology prediction tools were employed for orthology inference. Finally, the predicted orthology relationships were clustered into orthologous groups by MCL algorithm.

### Orthology Inference

OrthoMCL and FastOrtho found more orthology relationships than SwiftOrtho and OrthAgogue. This is because OrthoMCL and FastOrtho use the negative log ratio of the e-value as the edge-weighting metric. The BLASTP program rounds E-value *<* 10^−180^ to 0. Consequently, for homolgs with an e-value *<* 10^−180^, OrthoMCL and FastOrtho treat them as the RBHs, overestimating the number of orthologs. An example showing the OrthoMCL and FastOrtho overestimation can be found in Table S4.

#### Computational resource use

OrthoMCL v2.0 used the most CPU time and real time because of the required I/O operations. The RAM usage of OrthoMCL was 3.45GB, at the same time, the generated intermediate file occupied *>*19 TB disk space. OrthAgogue was the most real time efficient because its ability to exploit a multi-core processor. However, the RAM usage of OrthAgogue was more than 100GB which exceeds most workstations and servers. The orthology inference module of FastOrtho was the most memory-efficient among all the tools and it is also fast. SwiftOrtho was the most CPU time efficient although its real time was twice as OrthAgogue. Because the orthology inference module of SwiftOrtho was written in pure Python, we retested it by using the PyPy interpreter, an alternate implementation of Python [48]. The results show that the real run time of SwiftOrtho was close to OrthAgogue’s (Table S5)

### Cluster Analysis

OrthoFinder identified the smallest number of orthologous groups. Other tools identified many more orthologous groups than OrthoFinder, ranging from 36,901 to 51,297. The APC algorithm find fewer clusters than the MCL algorithm.

### Evaluation of Predicted Orthologous Groups

The quality of predicted orthologous groups is shown in Figure 3. OrthoFinder has the best recall, while SwiftOrtho and OrthAgogue have top precision values but lower recall values than other tools. Since SwiftOrtho and OrthAgogue use a more stringent standard to perform orthology inference, this strategy often increases precision but decreases recall [10, 20, 21].

Because SwiftOrtho uses its built-in homology search module and its recall is lower than BLASTP’s, this may also cause a reduction in the recall of orthologous groups. To eliminate this possibility, we made two replacements. We replaced SwiftOrtho’s homology module with BLASTP for SwiftOrtho and replaced BLASTP with SwiftOrtho’s homology module for OrthoMCL, FastOrtho, OrthAgogue, and OrthoFinder. We then reran the orthology analysis on *Euk*. The results show that for most tools replacing BLASTP with SwiftOrtho’s built-in homology search module does not significantly reduce the recall (Figure 4). The difference in recall between using SwiftOrtho’s homology search and using BLASTP is less than 3% except for OrthoMCL and FastOrtho. The recall for OrthoMCL and FastOrtho decreased by 5% and 8%, respectively. The most likely reason is that the E-value of SwiftOrtho’s homology search module is more precise than that of BLASTP, which reduces the false RBHs as mentioned above. These results also show that SwiftOrtho’s homology search module is a reliable and fast alternative to BLASTP.

Since SwiftOrtho uses an APC clustering algorithm, we ran SwiftOrtho with MCL and APC on the same data. The results (Figure 5) show that performance of APC is very close to that of MCL. APC improves the recall of most tools (Figure 5). These results also show that APC is a reliable alternative to MCL. APC requires less memory and can be used to cluster large-scale data.

**Figure 5.**
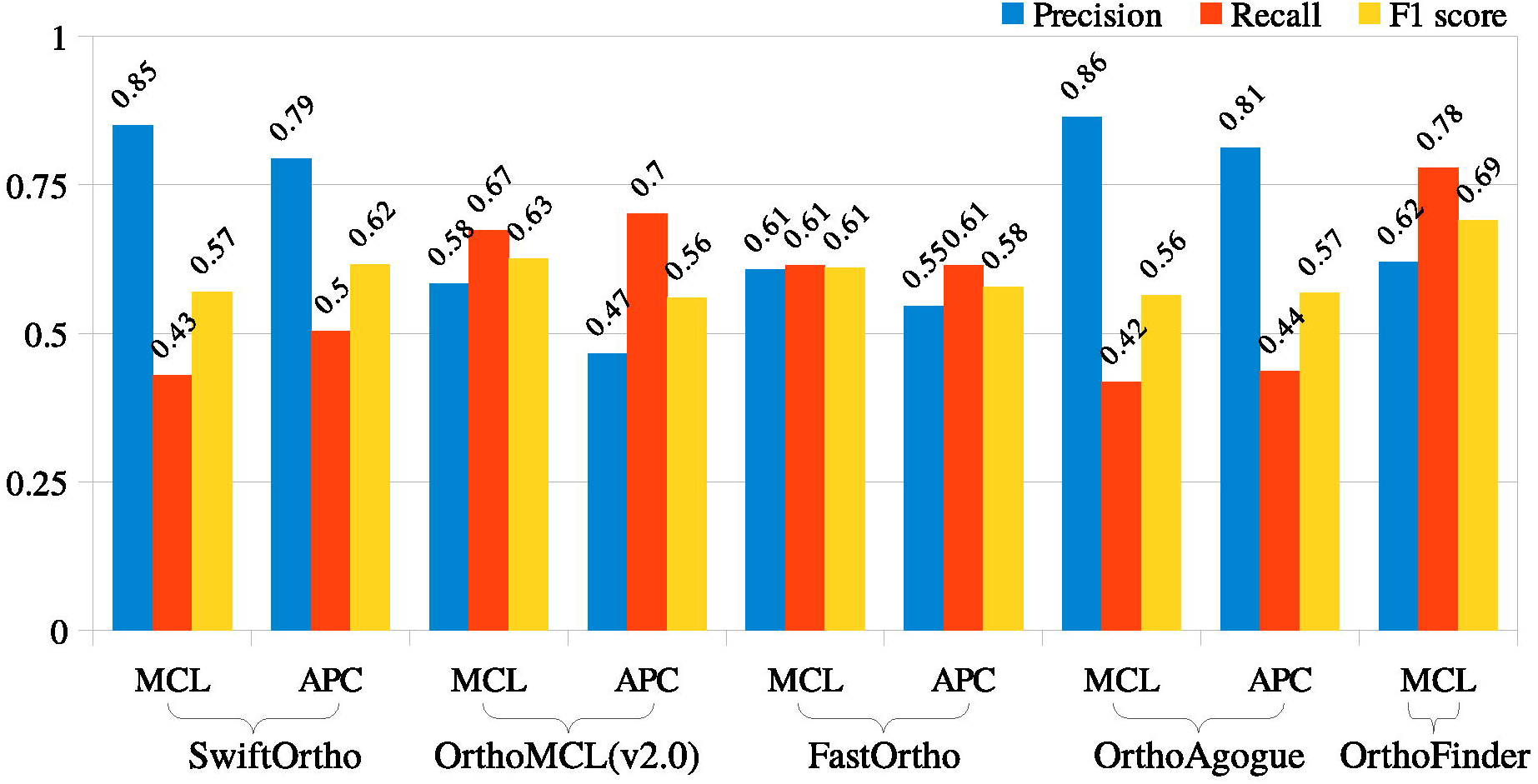
Markov Clustering versus Affinity Propagation Clustering. Both algorithms were applied to cluster the orthology relationships of the Euck set inferred by different orthology prediction tools, into orthologous groups. As OrthFinder does not report orthology relationships, the Affinity Propagation can not apply to its results. **MCL:** Markov Clustering algorithm; **APC:** Affinity Propagation Clustering.

### Orthology Analysis on *QfO 2011*

The results of the orthology analysis on *QfO 2011* are shown in Table 2:

**Table 2.**
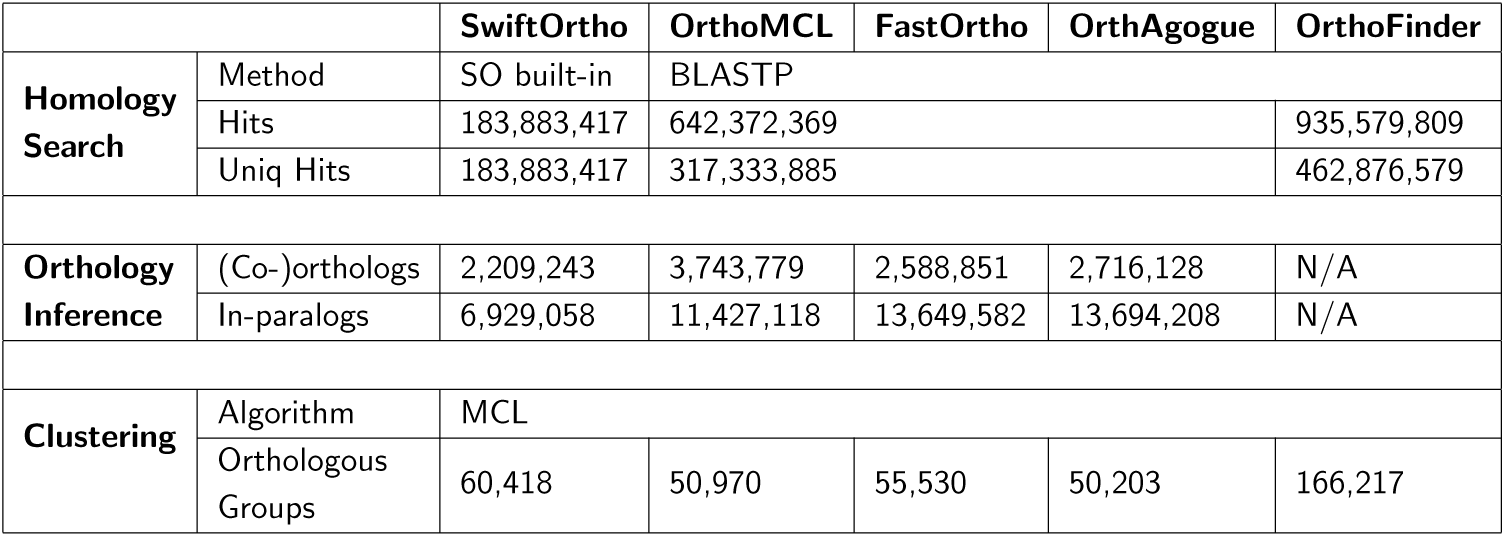
Comparative orthology analysis on the Quest for Orthologs reference proteome 2011 dataset. SO: SwiftOrtho; MCL: Markov Clustering; APC: Affinity Propagation Cluster; N/A: not available.

### Homology Search

SwiftOrtho found 183,883,417 unique hits while BLASTP found 462,876,579 unique hits. However, SwiftOrtho is about 163 times faster than BLASTP.

### Orthology Inference

OrthoMCL found many more orthologs and co-orthologs than the other tools. SwiftOrtho found fewer in-paralogs than other available tools. The CPU time of SwiftOrtho is the least of all tools. When using the PyPy interpreter, the real time of SwiftOrtho is also close to that of the fastest one, OrthAgogue (Supplementary Table S6).

### Cluster Analysis

Overall, the clustering numbers of SwiftOrtho, OrthoMCL, FastOrtho, and OrthAgogue are similar.However, the number of clusters found by OrthoFinder is three times that of other tools, and the next evaluation also shows that OrthoFinder performed poorly on *QfO 2011*.

### Evaluation of Predicted Ortholog Relationships

The evaluation shows that the performance of SwiftOrtho is close to that of Inparanoid (Figure 6). In some tests (Figure 6, D-E), SwiftOrtho outperformed Inparanoid. SwiftOrtho had the best performance in the Generalized Species Tree Discordance Benchmark and Agreement with Reference Gene Phylogenies: TreeFam-A tests. In the Species Tree Discordance Benchmark, SwiftOrtho had the minimum Robinson-Foulds distance. In the Enzyme Classification (EC) conservation test, SwiftOrtho had the maximum Schlicker similarity. These two metrics reflect the performance of the algorithm in accuracy and the results show that SwiftOrtho has an overall higher accuracy than the other tools, at the same time, the recall of SwiftOrtho was lower in some of the QfO tests. The most probable reason is that when we performed all-*vs*-all homology search, we used a long seed which resulted in fewer homologs being detected.

**Figure 6.**
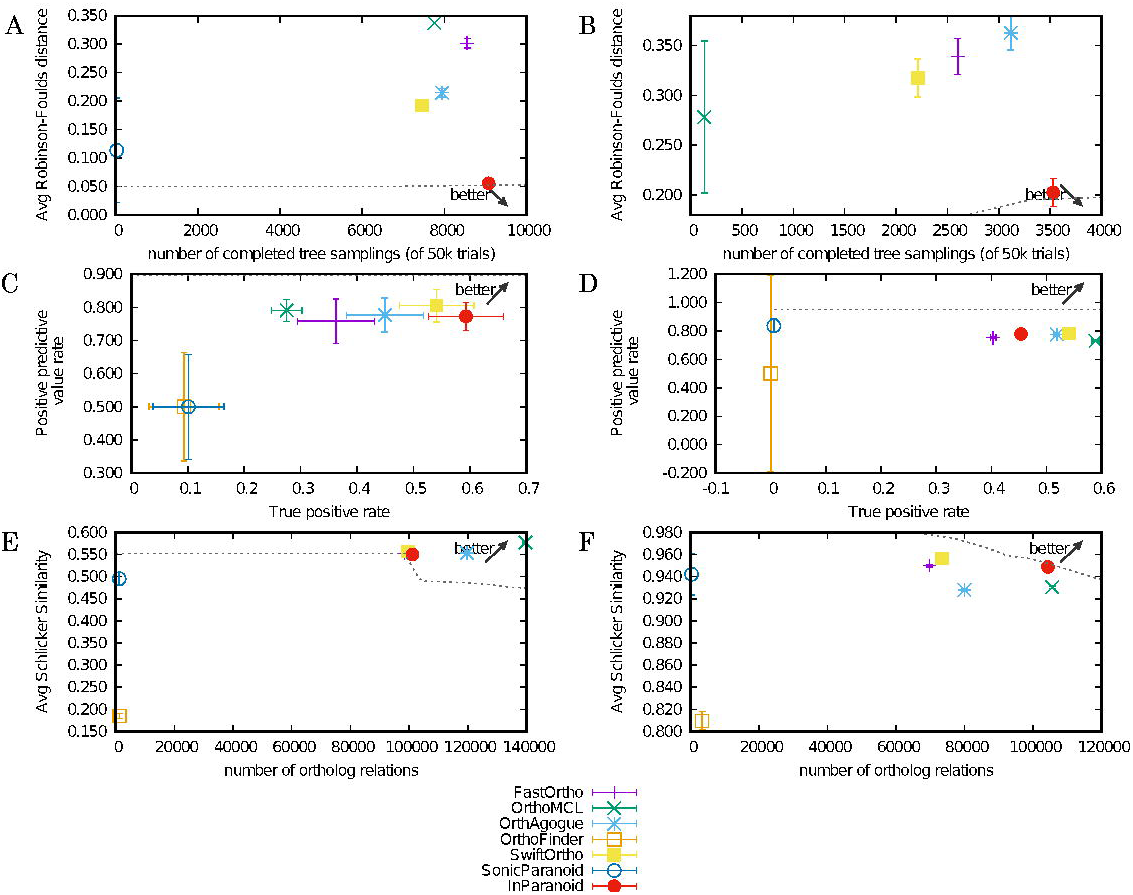
The Benchmarking in Quest for Orthologs. **A:** Species Tree Discordance Benchmark. Inparanoid has minimum average Robinson-Foulds distance. SwiftOrtho’s average RF distance is close to Inparanoid. The prediction inferred by OrthoFinder is not aviable in this test; **B:** Generalized Species Tree Discordance Benchmark. InParanoid has minimum average Robinson-Foulds distance. The prediction inferred by OrthoFinder is not aviable in this test; **C:** Agreement with Reference Gene Phylogenies of SwissTree. SwiftOrtho has the highest positive prediction value rate(Recall). InParanoid has the highest true positive rate(Precision); **D:** Agreement with Reference Gene Phylogenies of TreeFam-A. SonicParanoid has the highest positive prediction value rate(Recall), however, its true positive rate(Precision) is close to zero. SwiftOrtho has the second highest Recall and Precision; **E:** Gene Ontology conservation test. OrthoMCL has the highest average Schlicker similarity; **F:** Enzyme Classification conservation test. SwiftOrtho has the highest average Schlicker similarity. OrthoMCL detected the most orthology relationships and has the highest Recall.

### Orthology Analysis On *Bac*

The results of orthology analysis on *Bac* are shown in Table 3:

**Table 3.** Comparative orthology analysis on the Bac set. SO: SwiftOrtho; MCL: Markov Clustering; APC: Affinity Propagation Cluster; N/A: not available.

|  |  | SwiftOrtho |  | OrthoMCL | FastOrtho | OrthAgogue | OrthoFinder |
| --- | --- | --- | --- | --- | --- | --- | --- |
| Homology Search | Method | SO built-in |  |  |  |  | N/A |
|  | Hits | 8,478,732,753 |  |  |  |  | N/A |
|  | Uniq Hits | 8,478,732,753 |  |  |  |  | N/A |
| Orthology Inference | (Co-)orthologs | 876,766,940 |  | N/A | 950,683,849 | N/A | N/A |
|  | In-paralogs | 622,292 |  | N/A | 663,052 | N/A | N/A |
| Clustering | Algorithm | MCL | APC | MCL |  |  |  |
|  | Orthologous Groups | 240,162 | 167,355 | N/A | 242,816 | N/A | N/A |

### Homology Search

SwiftOrtho detected 8,966,131,536 homologs on the *Bac* set within 1,247 CPU hours. Because it takes long time to perform all-*vs*-all BLASTP search on the full *Bac*, we randomly selected 1,000 protein sequences from *Bac* and searched them against the full *Bac* set. It took BLASTP 5.1 CPU hours to find the homologs of these 1,000 protein sequences. We infer that the estimated CPU time of BLASTP on the full *Bac* set should be around 30,000 CPU hours. SwiftOrtho was almost 25 times faster than BLASTP on *Bac*.

### Orthology Inference

SwiftOrtho, OrthoMCL, FastOrtho, and OrthAgogue were used to infer (co-)orthologs and in-paralogs from the homologs detected by the homology search module of SwiftOrtho in the *Bac* set. We did not test Orthofinder, because Orthofinder does not accept a single file of homologs as input. For the 1,760 genomes in *Bac*, OrthoFinder needs to perform 3,097,600 pairwise genome comparisons, which will generate the same number of files. Then, OrthoFinder performs the orthology inference on these 3,097,600 files. Even at one minute per file, it will take an estimated six CPU years to process all the files.

Due to memory limitation, only SwiftOrtho and FastOrtho finished the orthology inference on *Bac*. The results are shown in Table 3. The numbers of (co-)orthologs and in-paralogs inferred by SwiftOrtho and FastOrtho are similar. The number of common orthology relationships between SwiftOrtho and FastOrtho was 861,619,519 (98.2% of SwiftOrtho and 90.57% of FastOrtho). Compared with *Euk*, SwiftOrtho and FastOrtho have a similar predictive quality on *Bac*. There are three possible explainations for these results. The first one is that *Euk* contains many protein isoforms which cause FastOrtho to overestimate the number of orthologs and in-paralogs. The second one is that the gene duplication rate in Bacteria is lower than that in Eukaryotes [49, 50]. For *Bac*, each gene in one species has only small number of homolgs in other species, which makes FastOrtho unlikely to overestimate the number of RBHs. The third one is that SwiftOrtho uses double-precision floating-point to store the E-value, which increases the precision of E-value from 10^−180^ to 10^−308^. This improvement also reduces the possibility that FastOrtho may report false RBHs.

#### Computational resource use

FastOrtho and OrthAgogue did not finish the tests due to insufficient RAM, OrthoMCL aborted after running out of disk space, as it needed more than 18TB. Only SwiftOrtho and FastOrtho finished the orthology inference step. The Peak RAM usage of SwiftOrtho and FastOrtho were 90.6GB and 99.5GB, respectively. When we used the PyPy interpreter, the Peak RAM usage of SwiftOrtho was reduced to 72.1GB. FastOrtho was about 1.52 times faster than SwiftOrtho which ran the tests in the CPython interpreter. When using the PyPy interpreter, SwiftOrtho ran 1.58 times faster than FastOrtho. The memory usage and CPU time are shown in Supplementary Table S7

### Cluster Analysis

The clustering numbers of SwiftOrtho and FastOrtho are similar. We compared the APC algorithm and the MCL algorithm, and APC found fewer clusters than MCL. The APC used much less memory and less CPU time than MCL. However, due to the lack of support for multi-threading and a large number of I/O operations, the real run time of APC is longer than that of MCL.

### Test on Low-memory System

Because SwiftOrtho is designed to handle large-scale data on low-memory computers, we used it to analyze *Bac* on a range of computers with different specifications. The results (Supplementary Table S8) show that the memory usage of SwiftOrtho is flexible and adaptes to the size of the computer’s memory. In the tests, SwiftOrtho finished an orthology anlaysis of *Bac* on computers with only 4GB RAM in a reasonable time (Table S8).

## Discussion

We present SwiftOrtho, a new high performance graph based homology classification tool. Unlike most tools that can only perform orthology inference, SwiftOrtho integrates all the modules necessary for orthology analysis, including homology search, orthology inference and cluster analysis. SwiftOrtho is designed to analyze large-scale genomic data on a normal desktop computer in a reasonable time. In our tests, SwiftOrtho’s homology search module was nearly 30 times faster than BLASTP. The orthology inference module of SwiftOrtho was nearly 500 times faster than OrthoMCL when applied to *Euk*. When applied to the large-scale dataset, *Bac*, SwiftOrtho was the only one that finished orthology inference test on a workstation with 32GB RAM. The cluster module of SwiftOrtho using APC can handle data that is much larger than the computer memory. In our test, APC has comparable recall and accuracy, but requires much less memory than MCL. APC even improved *F*_1_-measure score by increasing recall in most cases. With the help of these optimized modules, SwiftOrtho has successfully finished an orthology analysis of 1,760 bacterial genomes on a machine with only 4GB RAM. SwiftOrtho is not only fast but also accurate, as showing the results produced when running on orthobench and QfO[12, 24].

## Conclusion

In summary, SwiftOrtho is a fast, accurate orthology prediction tool that can analyze a large number of sequences with minimal computational resource use. The installation and configuration of SwiftOrtho is simple and does not require the user to have any experience in database configuration. It is easy to use, the only input required by SwiftOrtho is a FASTA format file of protein sequences with taxonomy information in the header line. Furthermore, SwiftOrtho is highly modular, and can be used to

SwiftOrtho can be integrated into various common pipelines where fast orthology classification is required such as pan-genome analysis, large-scale phylogenetic tree construction, and other multi-genome analyses. It is specifically suited for microbial community analyses, where large number of sequences and species are involved.

## Availability of data and materials

SwiftOrtho was written in Python 2.7 and is available at https://github.com/Rinoahu/SwiftOrtho under a GPLv3 license.

## Author’s contributions

Text for this section …

## Acknowledgements

Text for this section …

