## Supplementary Material for "SwiftOrtho: a Fast, Memory-Efficient, Multiple Genome Orthology Classifier"

### Methods

#### 1 Data sets and online service used in benchmarking

| Data or Service | URL |
|---|---|
| OrthoBench | http://eggnog.embl.de/orthobench2/orthobench2.all.data.tar.gz |
| QfO 2011 | ftp://ftp.ebi.ac.uk/pub/databases/reference_proteomes/previous_releases/qfo_release-2011_04/2011_04_reference_proteomes.tar.gz |
| QfO service | http://orthology.benchmarkservice.org/cgi-bin/gateway.pl |

Table S1: Data sets and online service used in benchmarking.

#### 2 Default reduced amino acid alphabet

| Categories | Amino acid |
|---|---|
| 1 | AST |
| 2 | CFILMVY |
| 3 | DN |
| 4 | EQ |
| 5 | G |
| 6 | H |
| 7 | KR |
| 8 | P |
| 9 | W |

Table S2: Default reduced amino acid table used by SwiftOrtho homology search module.

#### 3 Optimization of Space Seed for SwiftOrtho

SpEED was used to find the optimized space seed and the command line is:

```
$ SpEED 8 2 .25 100
```

#### 4 Orthology Analysis

##### 4.1 Software

software used in orthology analysis is listed in Table S3

| Name | Version | Source |
|---|---|---|
| BLASTP | 2.2.27+ | ftp://ftp.ncbi.nlm.nih.gov/blast/executables/rmblast/2.2.28/ |
| OrthoMCL | 2.0 | https://github.com/stajichlab/OrthoMCL |
| FastOrtho | 1.1.10 | https://github.com/davidemms/OrthoFinder/releases |
| OrthAgogue | | https://github.com/guyleonard/orthagogue |
| InParanoid | 4.1 | http://software.sbc.su.se/cgi-bin/request.cgi?project=inparanoid |
| SonicParanoid | 1.0 | http://iwasakilab.bs.s.u-tokyo.ac.jp/sonicparanoid/ |
| MCL | v14-137 | https://www.micans.org/mcl/index.html?sec_software |

Table S3: Software used in orthology analysis

#### 4.2 Parameters for running the software

The Orthology Analysis Pipeline for SwiftOrtho, OrthoMCL, FastOrtho, and OrthAgogue has three steps of homlogy search, orthology inference, and cluster analysis.

##### 4.2.1 Homology Search

1. SwiftOrtho for *Euk*:

```
$ python SwiftOrtho/bin/find_hit.py -p blastp -i input \
-d input -o output -r aa20 -e 1e-5 -s 1011111,11111
```

2. SwiftOrtho for *QfO 2011*:

```
$python SwiftOrtho/bin/find_hit.py -p blastp -i input \
-d input -o output -r aa20 -e 1e-5 -s 11111111
```

3. SwiftOrtho for *Bac*:

```
$python SwiftOrtho/bin/find_hit.py -p blastp -i input \
-d input -o output -r aa20 -e 1e-5 -s 111111
```

4. BLASTP for *Euk* and *QfO 2011*:

```
$blastall -p blastp -i input -d input -o output -m 8 \
 -v 1000000 -b1000000 -e 1e-5
```

##### 4.2.2 Orthology Inference

1. SwiftOrtho:

```
$python SwiftOrtho/bin/find_orth.py -i input \
 -c 0.5 -y 0 > output
```

2. OrthoMCL:

   We followed the instructions at the URL: `http://orthomcl.org/common/downloads/software/v2.0/Us`

3. FastOrtho:

```
$FastOrtho --option_file option_file.txt
```

4. OrthAgogue:

```
$orthAgogue -i input -e 5 -O output -b -c 8 \
 -dbs 1000000000 -u -A
```

###### 4.2.3 Cluster Analysis

1. Affinity propagation algorithm:

```
$python SwiftOrtho/bin/find_clpy -i input -d 0.5 \
-a apc > output
```

2. Markov Cluster algorithm:

```
$mcl input --abc -I 1.5 -o output -te 12
```

##### 4.3 Inparanoid, sonicparanoid, and OrthoFinder

###### 4.3.1 OrthoFinder:

```
$mcl input --abc -I 1.5 -o output -te 12
```

###### 4.3.2 OrthoFinder:

```
$orthofinder -f input_dir -a 12 -t 12
```

###### 4.3.3 SonicParanoid:

```
$sonicparanoid.py -i input_dir -o output_dir -t 8 -m fast
```

###### 4.3.4 InParanoid:

```
$inparanoid.pl fasta_file_A fasta_file_B > output
```

### Results

| Query(*Rattus norvegicus*) | Target(*Tetraodon nigroviridis*) | E-value | Bit score |
|---|---|---|---|
| **ENSRNOP00000063891** | **ENSTNIP00000002888** | 0 | 764(763) |
| ENSRNOP00000063891 | ENSTNIP00000013692 | 0 | 761(751) |
| ENSRNOP00000063891 | ENSTNIP00000006654 | 0 | 735(736) |
| ENSRNOP00000063891 | ENSTNIP00000009027 | 0 | 713(713) |
| ENSRNOP00000063891 | ENSTNIP00000002673 | 0 | 700(691) |
| ENSRNOP00000063891 | ENSTNIP00000009026 | 0 | 687(689) |
| ENSRNOP00000063891 | ENSTNIP00000022868 | 0 | 673(701) |
| ENSRNOP00000063891 | ENSTNIP00000001850 | 0 | 662(670) |
| ENSRNOP00000063891 | ENSTNIP00000009028 | 0 | 658(686) |
| ENSRNOP00000063891 | ENSTNIP00000013693 | 0 | 622(622) |
| ENSRNOP00000063891 | ENSTNIP00000002030 | 0 | 611(625) |
| ENSRNOP00000063891 | ENSTNIP00000009029 | 0 | 598(612) |
| ENSRNOP00000063891 | ENSTNIP00000009813 | 0 | 585(585) |
| ENSRNOP00000063891 | ENSTNIP00000015340 | 0 | 577(568) |
| ENSRNOP00000063891 | ENSTNIP00000003103 | 0 | 569(569) |
| ENSRNOP00000063891 | ENSTNIP00000005199 | 0 | 565(567) |
| ENSRNOP00000063891 | ENSTNIP00000008595 | 0 | 560(560) |

Table S4: An example of OrthoMCL and FastOrtho overestimating orthologs. The values in parentheses are the bit scores of reciprocal BLAST hits. The protein ENSRNOP00000063891 in *Rattus norvegicus* has 17 hits in *Tetraodon nigroviridis*, E-value of which are an e-value of zero. Among the 17 hits, ENSTNIP00000002888 has the highest bit score. According to the Reciprocal Best Hits (RBHs) rule, only ENSRNOP00000063891 and ENSTNIP00000002888 are true orthologs because of the highest bit score. However, OrthoMCL and FastOrtho treat all the 17 hits as RBHs because they have the same E-value of effective zero. SwiftOrtho and OrthAgogue can distinguish false RBHs because they use the bit score as edge-weighting metric. However, OrthoMCL and FastOrtho treat other hits as orthologs as they have the same zero E-value.

| | | **SwiftOrtho** | **OrthoMCL** | **FastOrtho** | **OrthAgogue** | **OrthoFinder** |
|---|---|---|---|---|---|---|
| **Homology Search** | Method | SO built-in | BLASTP | | | |
| | CPU Time(hr) | 18.7 | 724 | | | 1,371 |
| **Orthology Inference** | CPU Time(min) | 23(10.7*) | 5,175 | 27 | 44 | 807 |
| | Real Time(min) | 23.7(11.6*) | 7,617 | 29 | 10 | 188 |
| | Peak RAM(GB) | 10.1(10.1*) | 3.4 | 2.9 | 105.0 | 6.3 |

Table S5: Performance comparison of all orthology analysis tools on *Euk*. *:Time and RAM usage under PyPy interpreter; SO: SwiftOrtho; MCL: Markov Clustering; APC: Affinity Propagation Cluster.

| | | **SwiftOrtho** | **OrthoMCL** | **FastOrtho** | **OrthAgogue** | **OrthoFinder** |
|---|---|---|---|---|---|---|
| **Homology Search** | Method | SO built-in | BLASTP | | | |
| | CPU Time(hr) | 45.88 | 2,995 | | | 7,500 |
| **Orthology Inference** | CPU Time(hr) | .42(.24*) | 3.08 | .59 | .50 | 13.55 |
| | Real Time(hr) | .43(.31*) | 6.8 | 1.78 | .29 | 3.25 |
| | Peak RAM(GB) | 13.98(13.98*) | 3.38 | 3.23 | 68.29 | 6.36 |

Table S6: Performance comparison of all orthology analysis tools on *QfO 2011*. SO: SwiftOrtho; MCL: Markov Clustering; APC: Affinity Propagation Cluster; *:Time and RAM usage under PyPy; N/A: not available.

| | | SwiftOrtho | | OrthoMCL | FastOrtho | OrthAgogue | OrthoFinder |
|---|---|---|---|---|---|---|---|
| **Homology Search** | Method | SO built-in | | BLASTP | | | N/A |
| | CPU Time(hr) | 1,247 | | ~30,000† | | | N/A |
| **Orthology Inference** | CPU Time(hr) | 41.6(17.7⋆) | | N/A | 27.6 | N/A | N/A |
| | Real Time(hr) | 43.5(19.5⋆) | | N/A | 27.7 | N/A | N/A |
| | Peak RAM(GB) | 90.6(72.1⋆) | | N/A | 99.5 | N/A | N/A |
| **Clustering** | Algorithm | MCL | APC | N/A | MCL | N/A | N/A |
| | CPU Time(hr) | 41.4 | 12.7 | N/A | 57.7 | N/A | N/A |
| | Real Time(hr) | 6 | 26 | N/A | 7 | N/A | N/A |
| | Peak RAM(GB) | 81.3 | 39.5 | N/A | 67.8 | N/A | N/A |

Table S7: Performance comparison of all orthology analysis tools on *Bac*. SO: SwiftOrtho; MCL: Markov Clustering; APC: Affinity Propagation Cluster; ⋆:Time and RAM usage under PyPy; †: estimated by a subset of protein sequences, N/A: not available.

| **Hardware** | | | | |
|---|---|---|---|---|
| **Platform** | **Virtual Machine** | **Desktop** | **Workstation** | **Server** |
| **CPU Model** | E3-1271 v3 | E5-1620 v4 | i7-6800K | E5-2640 v3 |
| **CPU Frequency(GHz)** | 3.6 | 3.5 | 3.4 | 2.6 |
| **CPU Cores** | 4 | 8 | 12 | 16 |
| **RAM(GB)** | 4 | 16 | 32 | 128 |
| **File System** | Local Disk | | | Network File System |
| **Orthology Inference** | | | | |
| **Input homologs** | 8,478,732,753 | | | |
| **(Co-)Orthologs** | 876,766,940 | | | |
| **In-Paralogs** | 622,292 | | | |
| **CPU Time (hrs)** | 37 | 37.6 | 36 | 41.6 |
| **Real Time(hrs)** | 69 | 46.5 | 41.7 | 43.5 |
| **Peak RAM (GB)** | 2.58 | 11 | 19.3 | 90.6 |
| **Clustering** | | | | |
| **Algorithm** | Affinity Propagation | | | |
| **Orthologous Groups** | 167,355 | | | |
| **CPU Time (hrs)** | 7.3 | 10 | 12.1 | 12.7 |
| **Real Time (hrs)** | 56.5 | 60.5 | 35.7 | 26 |
| **Peak RAM (GB)** | 3.6 | 14.8 | 25 | 39.5 |

Table S8: Orthology Analysis of SwiftOrtho on *Bac* running on systems of different specifications. SwiftOrtho can even perform orthology inference and cluster analysis on a machine with only 4GB RAM. The peak memory usage of SwiftOrtho is scalable, which guarantees that it will run even on low-memory system.
